# Terminator-dependent facilitated recycling of RNA polymerase III couples transcriptional activation and chromatin remodeling in vivo

**DOI:** 10.1101/2023.02.28.530554

**Authors:** Vinesh Vinayachandran, Sushma Shivaswamy, Ashutosh Shukla, Nikitha Kendyala, Purnima Bhargava

## Abstract

Termination is a crucial step in generating the functional transcriptome of a cell. The short genes transcribed by RNA polymerase (pol) III are mostly found in the highly transcribed genomic loci. The mechanism responsible for their high transcription rate in vivo is not yet established. Transcription terminator-dependent facilitated recycling of pol III on naked DNA templates is reported to increase transcriptional output in vitro. We found that apart from defining the 3’-end of transcript, the transcription terminator is essential for achieving high-level chromatin transcription by pol III in vitro and in vivo. Using terminator-deficient *SNR6* gene templates or a recycling deficient pol III mutant, we show that the TFIIIC-dependent transcriptional activation of chromatin is a process with three closely linked components, viz. anti-repression to naked DNA levels with TFIIIC binding, TFIIIC-dependent chromatin remodeling for better accessibility of the sequence elements and terminator-directed full transcriptional activation. Measurement of pol III occupancy on different gene regions demonstrated a direct link between the high transcription rate and the terminator dependent recycling of pol III in vivo. This novel regulatory mechanism may be generally applicable to the highly transcribed genes in any cell and even for cancer management wherein pol III transcription is found highly elevated.

## Introduction

The transcription termination is a multi-step complex process (Richard and Manley 2009; Kuehner et al. 2011). All three RNA polymerases (Pols) follow different strategies but pausing of the polymerases near the terminator is a pre-requisite for termination by all (Richard and Manley 2009). In case of PoI II, there is a close correlation between elongation, termination and processing of the mRNA (Proudfoot et al. 2002; Moore and Proudfoot 2009; Mischo and Proudfoot 2013). Pol I requires proper cis- and trans-acting factors for proper termination and release of the transcript (Reeder et al. 1999) and shares some features with Pol II termination (El Hage et al. 2008; Kawauchi et al. 2008). The Pol III transcription generally terminates with the recognition of a specific T-stretch and requires factors like La, PC4 and NF1 (Cozzarelli et al. 1983; Maraia 1996; Wang et al. 2000; Arimbasseri et al. 2013). Yeast Pol III recognizes and pauses at its terminator sequence, an oligo dT cluster of six or more ‘T’ residues in the non-transcribed strand, in the absence of any other factor (Allison and Hall 1985; Braglia et al. 2005).

The highly conserved pol III transcription machinery constitutes the 17-subunit polymerase and three basal factors, TFIIIA, B and C, depending on the class of the gene transcribed in yeast (Acker et al. 2013; Arimbasseri et al. 2013). Target genes that characteristically have only intra-genic promoter, encode various short, non-coding RNAs required for various physiological functions (Dieci et al. 2007). Binding of TFIIIC to the intra-genic boxes A and B recruits the initiation factor TFIIIB at -30 bp position in the upstream gene region. Yeast *SNR6* (coding for U6 snRNA) gene is unusual in having its box B downstream of terminator and in not having TFIIIC requirement for naked DNA transcription in vitro since TFIIIB can bind to a TATA box present at the -30 base pair (bp) position (Margottin et al. 1991).

*SNR6* gene transcription in vivo and on chromatin templates assembled in vitro is dependent on the presence of TFIIIC and the extragenic box B (Brow and Guthrie 1990; Burnol et al. 1993a; Shivaswamy et al. 2004). The TFIIIC-induced chromatin remodeling results in the positioning of one nucleosome between its intragenic box A and extragenic box B with the gene terminator placed near the dyad axis (Shivaswamy et al. 2004). The resultant specific nucleosome positionings which were found essential for the transcriptional activation to more than naked DNA levels on the pol III-transcribed *SNR6* gene (Shivaswamy et al. 2004; Shivaswamy and Bhargava 2006) suggested a role for the terminator in the high level chromatin transcription.

Transcription termination depends on the nascent RNA displacement, required for polymerase release (Campbell and Setzer 1992). Recent in vitro studies on the pol III transcription termination have revealed the details of constituent steps and shown the role of terminator even in the RNA cleavage and release of the transcript and pol III (Arimbasseri and Maraia 2013; Arimbasseri and Maraia 2015; Mishra and Maraia 2019; Mishra et al. 2021). Re-initiation of transcription is reported as coupled to the termination process in vitro (Dieci and Sentenac 1996; Ferrari et al. 2004; Lyke-Andersen et al. 2011; Dieci et al. 2013). The preferential and terminator-facilitated pol III recycling on the same template allows new cycles to be more rapid than the initial one (Dieci and Sentenac 2003; Dieci et al. 2013). It is not yet ascertained whether the facilitated pol III recycling-reinitiation is employed by the genes in the highly transcribed genomic loci to meet the high cellular demand of the pol III transcripts in vivo. Though finding direct answer may be technically difficult in vivo, in vitro assays may be powerful tools to address this question.

Using a combination of in vitro and in vivo approaches, we found the transcription terminator plays more than one role in transcriptional activation on the chromatin. While it is required to define the physical end of transcribed region, and required for the facilitated recycling on naked DNA as previously reported (Dieci and Sentenac 1996); it couples the chromatin remodeling to transcription activation without influencing the nucleosome position as outcome of the remodeling process. The terminator, TFIIIC and ATP-dependent chromatin remodeling complement and cooperate with each other to facilitate the recycling of pol III, for achieving the full transcriptional activation of the pol III-transcribed *SNR6* chromatin. Our results provide direct evidences for the terminator-dependent pol III recycling in vivo, demonstrated so far only in vitro.

## Results

### Terminator influences the transcription efficiency of pol III

Native as well as modified *SNR6* gene constructs with variously located terminator were used to find its role in TFIIIC-dependent transcriptional activation of repressed *SNR6* chromatin. Measurement of naked DNA transcription by UTP-incorporation assay (Supplemental Fig. S1B) or primer extension method (Fig. 1B) in vitro shows similar levels from pU6LNST and pCS6, while pU6LNS and pU6LR are transcribed at comparatively lower level (Supplemental Fig. S1C). As compared to the pCS6 DNA, the absence of terminator at its normal position reduces transcription of pU6LNS and pU6LR (Fig. 1A) ∼3.5 fold. Linearization of a U6 gene carrying plasmid at a location upstream of the terminator was reported to give similar reduction (3 fold) of transcription compared to the yield from the U6 templates linearized downstream to the terminator (Dieci and Sentenac 1996).

**Figure 1.**
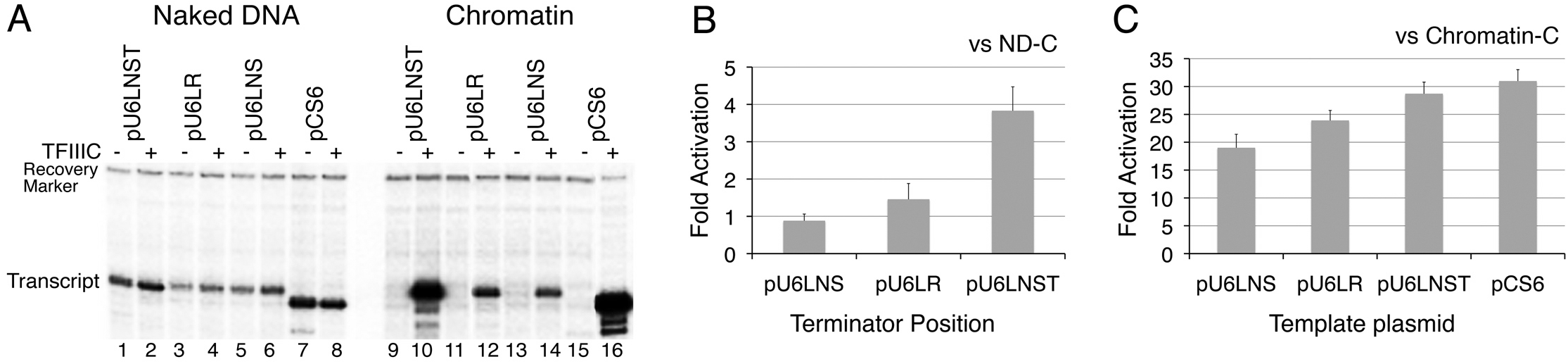
Terminator plays an important role in transcription activation on chromatin templates. (A) Requirement of terminator for transcriptional activation of modified U6 snRNA genes. A representative gel showing primer extension products of the transcripts (98-nt from pCS6 and 101-nt from the pU6LNS based plasmids) from the four plasmid DNAs is given. Positions of the recovery marker and the transcript are marked. (B) and (C) Transcriptional activation of chromatin templates with variously placed terminator due to TFIIIC. Fold activation gives increase of transcript yields (normalized with recovery marker) from chromatin in presence of TFIIIC with respect to (B) naked DNA (ND) and (C) chromatin in the absence of TFIIIC. Average from three independent experiments with scatter is shown.

Similar to pCS6, transcription of modified U6 naked DNA templates using highly purified components does not require TFIIIC (Fig, 1A). Chromatin formation results in complete repression, which could be abolished by TFIIIC addition (cf. chromatin lanes without or with TFIIIC, Fig. 1A). The transcript yield from pU6LNST chromatin is highest among the modified genes (Fig. 1A,B). When compared to respective naked DNAs, fold activation from chromatin in the presence of TFIIIC for pU6LR is only 1.5 fold higher than that from pU6LNS (Fig. 1B), while pU6LNST shows a 4 fold activation, similar to 5-6 fold activation from pCS6, reported earlier (Shivaswamy et al. 2004). With TFIIIC addition, repressed pU6LNS chromatin showed ∼20 fold (as opposed to ∼30 fold for pCS6 and pU6LNST) activation (Fig. 1C). Presence of terminator at a further downstream location in pU6LR reduces this activation to ∼10 fold (Fig. 1C). Naked DNA transcription from pU6LR is higher than pU6LNS (Supplemental Fig. S1C), but fold activation with respect to chromatin without TFIIIC is comparatively less (Fig. 1C), suggesting the terminator at abnormal position does not support TFIIIC-dependent activation from chromatin while high level of transcription requires a terminator at its normal location (Fig. 1B,C).

### Terminator does not direct nucleosome position on the *SNR6* gene

The terminator sequence in yeast *SNR6* is a stretch of 10 T:A base pairs, which generally behaves like a stiff rod (Nelson et al. 1987; Segal and Widom 2009). The least bent configuration of nucleosomal DNA at the dyad axis (Luger et al. 1997) raises the possibility of a role for the terminator in positioning of the nucleosome between the boxes A and B. As chromatin structure on a gene may influence its transcription, we checked for the possibility of a different chromatin structure on the modified *SNR6* gene constructs pU6LNS and pU6LR (Supplemental Fig. S1A, Table S1).

Similar to the wild type gene in pCS6 (Shivaswamy et al. 2004), chromatin structure analysis by the indirect end labeling (IEL) found a TFIIIC-dependent nucleosome positioning between the boxes A and B (Fig. 2A) on the modified genes. The micrococcal nuclease (MNase) digestion pattern of chromatin and naked DNA without or with TFIIIC were indistinguishable (Fig. 2A), indicating no preferred positioning of nucleosomes in the gene region. With TFIIIC addition, a protected segment of ∼200 bp size is seen between the boxes A and B (black ellipse) as well as downstream of box B (gray ellipse). Comparison of the digestion profiles of similarly digested chromatin (Supplemental Fig. S2) confirmed this nucleosomal size protecion for both the DNAs. High-resolution MNase footprinting confirmed similar binding of TFIIIC and no difference between the exact positions of the nucleosomes on both the plasmids (Fig. 2B,C,D). Profile comparison of the chromatin with/without TFIIIC (Fig. 2C) maps the span of nucleosomal protection between the bp +42 and +182 (dyad at +112 bp) on both the plasmids; which is similar to the protection between bp +50 to +190 (dyad at +120 bp) in case of pCS6 (Shivaswamy et al. 2004). An upstream shift of ∼8 bp in position of the nucleosomes could be because of a distance reduction between the boxes A and B by ∼17 bp in the modified genes. This analysis rules out a role for the terminator in nucleosome positioning but implicates the terminator role in the TFIIIC-dependent activation process from the chromatin.

**Figure 2.**
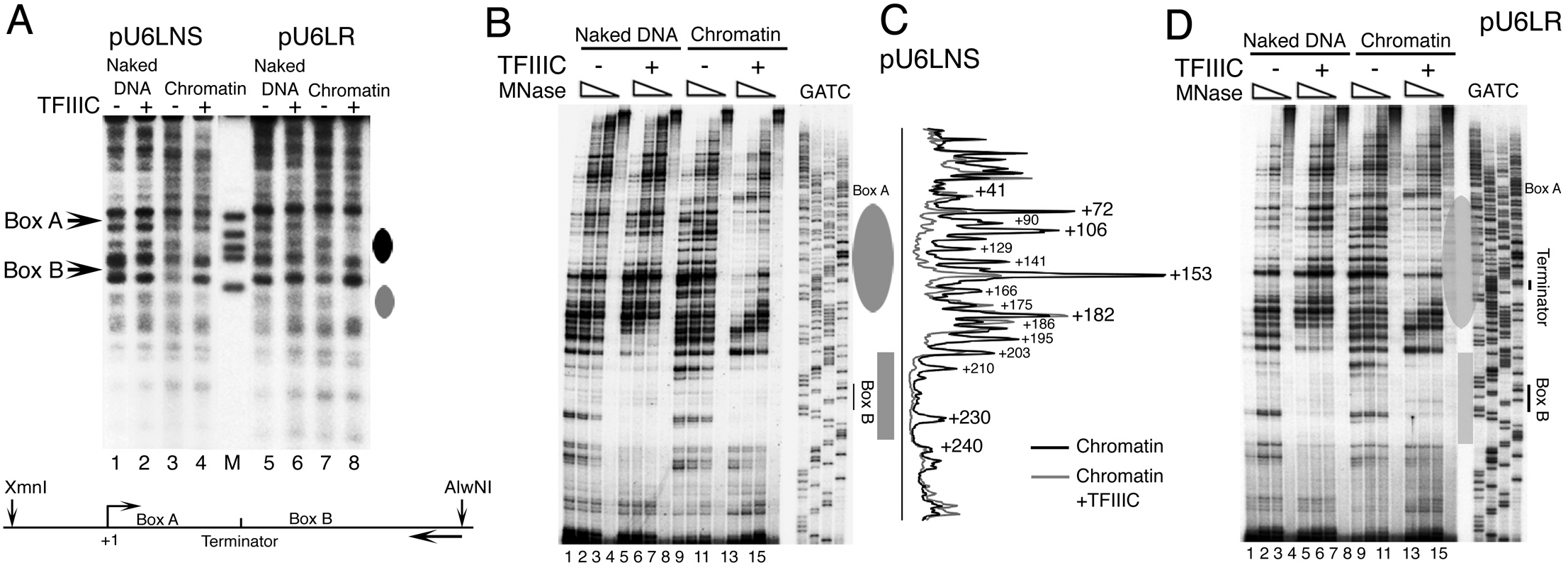
Chromatin Remodeling is independent of transcription terminator. Chromatin structure analysis of the modified U6 genes without (pU6LNS) or misplaced (pU6LR) terminator shows a positioned nucleosome between the boxes A and B in the presence of TFIIIC. Chromatin was assembled on plasmid DNAs and incubated with saturating amounts (11 fold molar excess) of TFIIIC where indicated for 30 mins., before digesting with MNase. (A) Indirect End-labeling (IEL) analysis. The Southern blots of MNase digestion products after secondary digestion with restriction enzymes AlwNl and Xmnl were probed with a primer that hybridizes to the gene proximal side of the AlwN 1 site, ∼900 bp downstream of the box B on both the plasmids (bottom cartoon). The autorad was calibrated using molecular weight markers loaded on the same gel (lane M) to find the size of the observed protections. Protection due to positioned nucleosomes in lanes 4 and 8 are marked by ellipses on the right hand side of the gel. (B) and (D) High resolution MNase footprinting of the modified U6 genes, pU6LNS (panels B,C) and pU6LR (panel D) without or with TFIIIC addition. Lanes 1-8 represent naked DNA while lanes 9-16 show digestion products of the chromatin in both the gels. In the each set of four lanes, first three lanes have samples subjected to three levels of MNase while the last one shows primer extension over the undigested sample. The primer hybridized 58 bp downstream of the box B, complementary to the top strand. Sequencing lanes under GATC were used to locate and mark the boxes A, B and terminator. The rectangle shows protection over the box B while the ellipse represents the positioned nucleosome in both the panels. (C) The profile matchings of lanes 11 (chromatin) and 15 (chromatin +TFIIIC) from the gel in panel B. Numbers denote the positions cleaved by the MNase on the gene. The peak at +230 represents the single cleavage in box B.

### Transcription terminator is required for chromatin activation

To confirm that the terminator at native location is responsible for the high level transcription from the *SNR6* chromatin; terminator sequence of the wild type gene in pCS6 was disrupted (rather than deleted; Fig. 3A, pU6dT). Naked pU6dT is transcribed at very low level (Supplemental Fig. S1) and anti-repression by TFIIIC can activate the repressed pU6dT chromatin only to naked DNA level (Fig. 3B lanes 1,2). This is similar to the results in Figure 1C where pU6LNS and pU6LR also are activated to only naked DNA levels, confirming the requirement of proper terminator for full activation of chromatin. Accordingly, in contrast to ∼30 fold activation of pCS6, ∼9 fold activation of pU6dT with respect to repressed chromatin (Fig. 3C) is comparable to ∼10 fold transcriptional activation of pCS6 chromatin with respect to naked DNA. These results indicate that chromatin formation represses the naked DNA transcription and terminator disruption influences the transcriptional activation.

**Figure 3.**
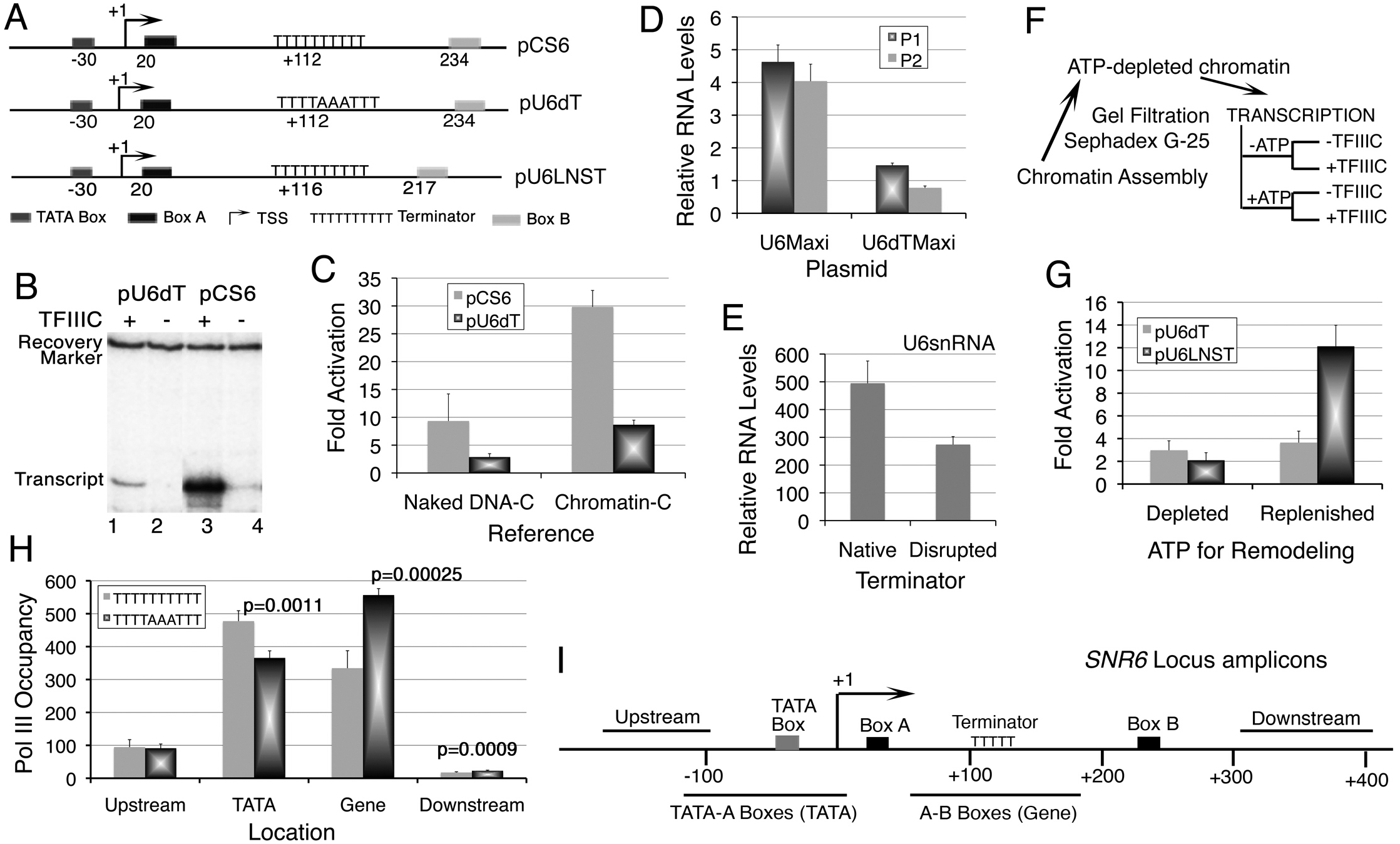
Transcription terminator facilitates pol III recycling on the yeast *SNR6* chromatin. (A) Schematic map of the *SNR6* gene in plasmids pCS6 (normal terminator), pU6dT (terminator disrupted in pCS6) and pU6LNST (terminator restored in pU6LNS). Numbers denote positions of the respective elements with respect to the TSS at +1. (B) In vitro transcription from the plasmids pU6dT and pCS6 assembled into chromatin. A representative gel showing primer extension products of the transcripts in the absence or presence of TFIIIC is shown. The recovery marker and transcript positions are marked. (C) Quantitative comparison of transcription from pU6dT and pCS6 chromatin. Fold activation of chromatin in presence of TFIIIC was measured for both the templates by calculating the increase with respect to transcription without TFIIIC from naked DNA (Naked DNA-C) or chromatin (Chromatin-C). Average from three independent experiments with scatter is shown. (D) Maxi RNA levels from yeast cells carrying the plasmids pU6Maxi or pU6dTMaxi, normalized to endogenous U6 snRNA levels. Average with scatter of measurements from at least three biological replicates is shown. (E) Quantification of U6 snRNA levels from the YPB17 and YPB110 cells carrying pRS413 plasmid with a wild type gene copy. Endogenous *SNR6* gene copy at its locus carries a functional native (wild type) or disrupted (mutant) terminator, indicated as their sequences in the legends of the graph. U6 levels were normalized against U4 RNA levels. Average from three independent biological replicates, with scatter is shown. (F) Scheme of ATP-depletion experiment. Chromatin assembled in the presence of ATP is gel filtered through Sephadex G-25 and equal aliquots are used for in vitro transcription without or with replenishing ATP, in the absence or presence of TFIIIC. (G) Requirement of ATP-dependent chromatin remodeling for transcriptional activation. The presence or absence of ATP refers not to transcription but to the larger amount of ATP (3 mM) required for the chromatin assembly and remodeling. Fold activation represents the ratio of normalized transcript yield in presence and absence of TFIIIC. Average from three independent experiments with scatter is shown. (H) Pol III (Rpc160-myc) occupancy on different regions of the *SNR6* gene locus in the cells carrying wild type or disrupted terminator was measured using ChIP-Real Time PCR method. Average from minimum three independent biological replicates, with scatter is shown. The p values of significance are given on the panel. (I) Positions of different amplicons on the *SNR6* gene locus, used for the Real Time PCR. Numbers denote the bp positions with reference to the TSS. Since gene copy in pRS413 plasmid is from only -120 to +312 bp of the genomic DNA, the upstream and downstream amplicons probe only the genomic locus. The TATA-A Boxes (TATA) and A-B Boxes (Gene) amplicons give total occupancies on both the genomic regions, native as well as the plasmid.

### Disruption of terminator results in loss of transcription from *SNR6* in vivo

Chromatin remodeling around the gene regulates the expression from the U6 gene locus in vivo (Arimbasseri and Bhargava 2008). Therefore, terminator-dependent coupling of chromatin remodeling and transcriptional activation may be a mechanism to generate high levels of transcript from this single copy gene in vivo as well. A maxi form of the gene in a plasmid with ∼59 bp insert in the transcribed region of the gene (Supplemental Fig. S3A,B) allows to differentiate the maxi U6 product from the normal U6 RNA in vivo (Burnol et al. 1993b). Similar to the in vitro data, with normalization against endogenous U6 RNA control, 3-4 fold lower yield from the U6dTMaxi (with disrupted terminator sequence) as compared to the U6Maxi transcript level (Fig. 3D), shows the terminator requirement for high transcript yield in vivo.

The chromatin structure around a gene may affect its expression. Due to the known differences in chromatin structure over the plasmid and genomic copy of the gene in vivo (Marsolier et al. 1995), we disrupted the gene terminator at its normal locus and rescued the cells from its deleterious effects by using a normal gene copy on the single copy plasmid pRS413 (Sikorski and Hieter 1989). The chromatin structure of the gene in situ did not show any difference between nucleosome positions at the *SNR6* locus in wild type or mutant cells (not shown). Measurement of total U6 levels in these cells (Fig. 3E) shows 50% lower U6 levels in absence of the functional terminator, affirming the functional terminator requirement to generate normal transcript levels in vivo.

### Transcriptional activation is coupled to chromatin remodeling

We had earlier reported that TFIIIC binding and ATP-dependent chromatin remodeling are two separable events underlying full transcriptional activation of the yeast *SNR6* chromatin (Shivaswamy and Bhargava 2006). Binding of TFIIIC to the gene makes the transcription by pol III through the gene body nucleosome possible, which is enhanced further 3 fold by the chromatin remodeling in the next stage (Shivaswamy et al. 2004). As the chromatin remodeling is unaffected, the different levels of transcriptional activation may be due to the presence or absence of a transcription terminator in the templates as reported earlier (Shivaswamy and Bhargava 2006). In order to confirm this possibility, transcription was followed as given under the scheme (Fig. 3F), with or without ATP-dependent chromatin remodeling (Shivaswamy et al. 2004). Transcription showed similar dependence on TFIIIC since despite ATP depletion, TFIIIC binding activates the transcription from both the chromatinized plasmids similarly (2-3 fold, Fig. 3G). With replenished ATP levels, transcription from pU6dT (terminator disrupted) chromatin does not increase significantly while similar to pCS6, a further ∼6 fold increase is seen from pU6LNST (terminator restored) chromatin. A loss of ∼3 fold activation on pU6dT chromatin is consistent when compared to either pCS6 (panel C) or pU6LNST (panel G) chromatin. The result shows that while anti-repression by TFIIIC binding restores naked DNA transcription level, further activation is linked to the chromatin remodeling and functional terminator.

### Terminator facilitates Pol III recycling on *SNR6* chromatin in vivo

Termination and re-initiation are two closely linked processes (Dieci et al. 2013). Therefore, reduced transcription with terminator disruption could be due to abrogation of the re-initiation. Recent studies have uncovered the in depth details of transcription termination by yeast pol III (Arimbasseri and Maraia 2015; Girbig et al. 2022; Xie et al. 2022). Pol III requires the T3, T4 (3^rd^, 4^th^T) residues of the terminator sequence in the non-template strand for conversion of the elongation complex into the metastable pre-termination complex (PTC). While T5 is required for the transcript release, its absence allows read-through with a pausing of the pol III, eventually releasing the transcript and pol III at T7 (Arimbasseri and Maraia 2015).

The terminator sequence in the plasmid pU6dT consists of an A at the 5^th^ position, which suggests a read-through of the terminator. A check for the downstream transcript from the pU6dT by primer (Supplemental Fig. S3C) extension on RNA from the in vitro transcription of naked pCS6 by the wild type pol III could barely detect a downstream transcription from pCS6 (Supplemental Fig. S3D). In contrast, equivalent transcription from the downstream as well as the gene region of pU6dT (Supplemental Fig. S3D lanes 3,4) suggests that in the absence of the 5^th^ T residue of the terminator, pol III continues transcription past the terminator without falling off the template. In the earlier study (Arimbasseri and Maraia 2015), the template used was a linear DNA with a very short continuation after the terminator sequence, such that being very close to the 3’ end of the template after the read-through to T7/T9 of the terminator, pol III falls off the template. With plasmid or genomic DNA as template in this study, pol III transcribes the region downstream of terminator but eventually falls off the template, releasing the transcript. Due to the dissociation of the transcription complex and termination of the ongoing cycle, pol III may not recycle-reinitiate on the same template culminating into reduced gene transcription when T_10_ stretch of the U6 terminator sequence is disrupted (Fig. 3D,E).

In order to visualize the loss of recycling in vivo, we measured the pol III occupancy on different regions of the *SNR6* gene (Arimbasseri and Bhargava 2008). As compared to the wild type, pol III occupancy on the *SNR6* gene carrying disrupted terminator (Fig. 3H) shows a loss at the TATA-A box initiation region and significant increase in the gene region including the DNA between the terminator and B box as well as the region downstream of B box (Fig. 3I). The increased occupancies show that pol III indeed goes past the disrupted terminator and pauses in the downstream region from where it falls off leading to its reduced levels near the transcription start site (TSS). Thus, the reduced pol III occupancy near the TSS, owing to its failure to recycle-reinitiate because of the disrupted terminator in vitro as well as in vivo directly results in the reduced transcript yield from *SNR6* (Fig. 3D,E).

### Transcriptional activation and terminator-dependent facilitated recycling of Pol III on chromatin are linked

Above results show that disruption of the transcription terminator, essential for efficient recycling of yeast pol III in vitro (Dieci and Sentenac 1996), disables the pol III recycling in vivo as well. Single round transcription assays used to find the recycling of pol III on the same template in vitro, involve heparin addition to trap the released enzyme molecules. This assay, the best available tool for naked DNA templates in vitro, faces a major drawback with chromatin, the physiological template in vivo. It has been known that heparin addition results in disruption of chromatin structure of the template (Courvalin and Dumontier 1982; Supplemental Fig. S4A), making subsequent transcription measurements wrong. Therefore, as an alternative, we utilized a mutant form of pol III (Pol IIIΔ) for in vitro transcription of pCS6 and pU6dT (Supplemental Fig. S4B). The mutant pol III shows proper initiation but does not recycle on naked DNA templates in vitro (Landrieux et al. 2006) owing to the absence of its three unique subunits C53, C37 and C11. The role of each subunit has been extensively investigated and it is established now that these subunits are crucial for several steps of termination including terminator recognition, PTC formation, RNA 3’ end cleavage, transcript release, pol III recycling and reinitiation on the same template etc. (Arimbasseri et al. 2013; Arimbasseri and Maraia 2015; Mishra et al. 2021). As Pol IIIΔ is lacking all these activities, the high transcriptional output by the wild type pol III could be directly linked to the end-effect, the terminator-dependent facilitated recycling-reinitiation.

Pol IIIΔ shows very little transcription from the naked pCS6 DNA without or with TFIIIC (Fig. 4A). As compared to the wild type, pol IIIΔ gives only naked DNA level transcription from pCS6 chromatin even in the presence of TFIIIC while no transcription could be seen from pU6dT (Supplemental Fig. S4B lanes 9,10,13,14). Strikingly similar (∼4 fold) activation levels with respect to the naked DNA level (Fig. 4C) from pCS6 (normal terminator) by pol IIIΔ and from pU6dT (disrupted terminator) by wild type pol III demonstrate the crucial link of the terminator with pol III recycling in the TFIIIC-dependent activation of chromatin transcription. Thus, terminator and pol III recycling, may be contributing to the transcriptional activation on the chromatin templates via a common path, the facilitated recycling of pol III.

**Figure 4.**
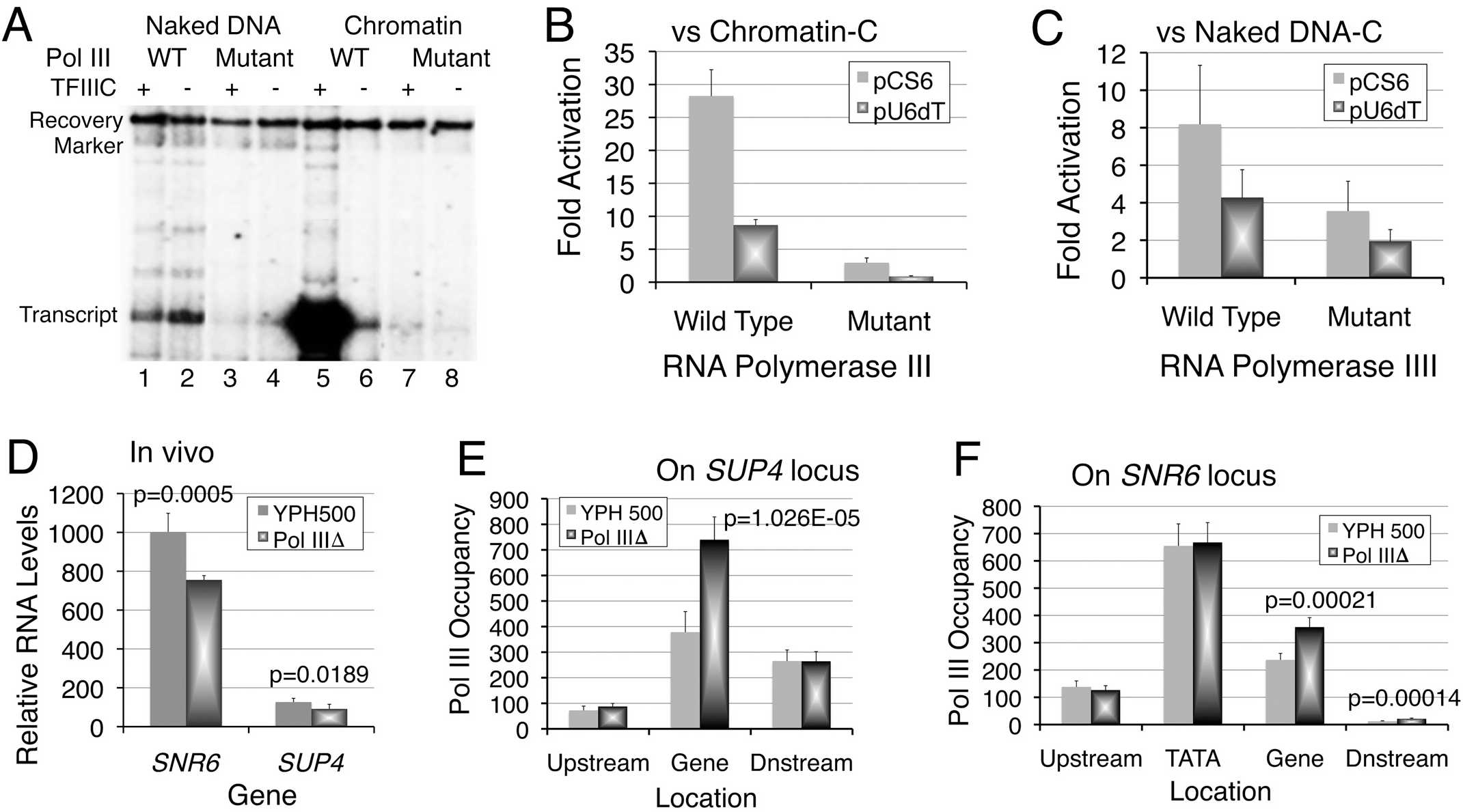
Terminator-dependent pol III recycling is required for transcriptional activation from chromatin. (A) In vitro transcription of pCS6 by wild type (WT) or mutant Pol 111 in the absence or presence of TFIIIC. A representative gel is shown. (B) and (C) Quantitative comparison of transcription by the wild type and mutant pol III from pCS6 and pU6dT chromatin. Fold activation represents the ratio of transcript yields from chromatin plus TFIIIC and chromatin (B) or Naked DNA (C) without TFIIIC. Average with scatter form three or more independent measurements is shown. (D) Primary transcript levels of *SNR6* and *SUP4* genes in the cells with wild type or mutant pol III show significant differences. Normalization was with U4 snRNA levels and average with scatter from at least three independent experiments is shown. The p values of significance are given. (E) and (F) Pol IIIΔ occupancy on the gene region in vivo is higher than the wild type pol III occupancy. Pol III (C128-Myc subunit) occupancy at the *SUP4* (E) and *SNR6* (F) gene loci. Upstream and Downstream represent the flanking regions of the gene in situ. Occupancy represents relative enrichment of wild type or mutant pol III over TEL VIR region. Graph shows average with scatter from at least three biological replicates. The p values of significance are given for the pairs.

### Terminator and chromatin regulate facilitated recycling of pol III in vivo

In order to validate the possibility of this mechanism operating in vivo, we used the cells carrying wild type or mutant enzyme for measuring the transcript levels and pol III occupancies on two pol III-transcribed genes *SNR6* and *SUP4* (tRNA^Tyr^). Both the genes were used earlier to demonstrate the terminator-dependent facilitated recycling of pol III in vitro (Dieci and Sentenac 1996) and shown to have influence of chromatin context on their expression in vivo (Arimbasseri and Bhargava 2008; Mahapatra et al. 2011). Significantly lower transcript levels from both *SNR6* and *SUP4* (tRNA^Tyr^) genes in the cells carrying pol IIIΔ (Fig. 4D) are consistent with the involvement of terminator-dependent pol III recycling in transcription at these loci in vivo.

As compared to in vitro, no specific transcript could be detected from the downstream region of the *SNR6* gene in vivo with either the wild type or the recycling-defective pol III (Supplemental Fig. S4C). This is consistent with the possibility of either the reported terminator arrest of pol IIIΔ (Arimbasseri and Maraia 2015) or the transcript degradation by the cellular surveillance machinery in vivo. In comparison of wild type, lower *SNR6* and *SUP4* transcript levels in cells carrying pol IIIΔ (Fig. 4D) suggest a lower occupancy of pol IIIΔ on these genes. We measured pol III enrichment on both the genes in both cell types by ChIP-Real Time qPCR method using primer pairs (Fig. 3I, Supplemental Fig. S4D) used earlier (Arimbasseri and Bhargava 2008; Mahapatra et al. 2011). Both the genes show similar distribution of pol III; higher occupancy in the gene region compared to the flanking regions (Fig. 4E,F). Consistent with normal initiation properties of pol IIIΔ (Landrieux et al. 2006), equal levels of both polymerases are found in the TATA-A box region of *SNR6* (Fig. 4E). Equal levels near the TSS and gene flanking regions but higher levels of pol IIIΔ on the transcribed region of both the genes (Fig. 4E,F) are consistent with the reported faster elongation rate and longer dwelling time near the terminator of the pol IIIΔ (Landrieux et al. 2006). Pol IIIΔ was shown to have termination as well as transcript release-reinitiation defect (Arimbasseri and Maraia 2013; Mishra et al. 2021). The terminator arrest and subsequent release of the pol IIIΔ (Arimbasseri and Maraia 2013) eventually lead to the highly reduced transcript yields (Fig. 4A). The absence of pol III from the region downstream of *SUP4* terminator in pol IIIΔ cells in vivo is consistent with its reported readthrough of the first oligo dT stretch but release at the second terminator of *SUP4* gene (Landrieux et al. 2006). Similarly, the absence of a downstream transcript (Supplemental Fig. S4C) but significantly higher pol IIIΔ occupancy on the *SNR6* locus could be due to its falling off the *SNR6* T_10_ terminator and not a readthrough followed by degradation of a downstream transcript in vivo. These measurements clearly show that with its all termination defects, the highly reduced transcriptional output of pol IIIΔ is because it fails to recycle in vivo.

Taking together, above results show that the terminator in its normal location is indispensable for the full transcriptional activation but not required for nucleosome positioning or chromatin remodeling. ATP-dependent chromatin remodeling or TFIIIC alone can give only partial activation of the transcription. Together, they make the terminator-dependent facilitated recycling of pol III possible on the chromatin, resulting in high level transcription of the *SNR6* gene. Pol III recycling defect or non functional terminator both lead to similar loss of transcription in vitro and in vivo. Monitoring pol III occupancy on different gene regions in vivo, we conclude it is the pol III recycling-reinitiation and not the termination alone, which is important and responsible for high transcriptional activation on chromatin templates in vivo. The in vivo occurrence of the pol III recycling also explains how the mobility of a single nucleosome in the downstream region of a large number of pol III-transcribed tRNA genes (including the *SUP4* gene) regulates their transcription earlier reported by us (Mahapatra et al. 2011; Kumar and Bhargava 2013; Shukla and Bhargava 2018), as discussed below.

## Discussion

The question whether terminator-dependent facilitated recycling of pol III takes place in vivo, has been a long-standing question. This study shows that the terminator dependent pol III recycling is a key requirement for high level transcription of the pol III-transcribed genes. Its operation in vivo on two different classes of the pol III-transcribed genes establishes high transcriptional rate and yield, in two apparently different but inherently similar ways (Supplemental Fig. S5). On both *SNR6* and *SUP4* genes, the transcriptional activation of the chromatin constitutes three major steps coupled to each other. (i) TFIIIC binding to the chromatin causes anti repression, which gives a naked DNA level transcription. (ii) Coupling of TFIIIC binding and anti-repression to ATP-dependent chromatin remodeling mobilizes the repressive nucleosomes to positions more conducive for transcription, resulting in a 3-4 fold increase. (iii) Transcription terminator links the further transcriptional activation to this chromatin remodeling via the terminator-dependent recycling of pol III. Based on these details, following discussion elaborates the applicability of the three steps to both the genes.

### Terminator-dependent facilitated pol III recycling on the yeast *SNR6* gene

The *SNR6* gene could be one of the few examples where a positioned nucleosome on the gene body facilitates its transcription (Shivaswamy et al. 2004). While the first two steps mentioned above were clearly demonstrated earlier as well (Shivaswamy et al. 2004; Shivaswamy and Bhargava 2006), this study has revealed the terminator dependent third step and its coupling to the other two steps. Apart from aligning the boxes A and B of the *SNR6* gene, wrapping of the gene body into a nucleosome brings the normally located terminator (in pCS6 or pU6LNST) near/at the dyad axis, close to the TSS of the gene in space. It is conceivable that juxtaposing the initiation region with the terminator near the dyad axis would facilitate direct transfer of pol III from terminator to initiation region and allow faster re-initiation on a nucleosome, giving transcription at higher than the naked DNA level. In contrast, this facilitatory effect on pol III re-cycling for the next round of transcription would not be possible from the wrongly placed terminator at +163 bp (∼50 bp downstream) in pU6LR, which goes to the other half of the nucleosomal DNA, away from the TSS. In both these arrangements, terminator is moved away from its potentially interfering location between the boxes A and B, enabling smooth TFIIIC binding and resulting in an anti repression to only naked DNA level transcription.

### Transcriptional activation of the chromatin and pol III recycling-reinitiation

As opposed to a positioned nucleosome on the *SNR6* gene body, a downstream nucleosome regulates transcription from the *SUP4* tRNA gene in vivo by influencing the accessibility of the gene terminator (Mahapatra et al. 2011). The yeast tRNA genes reside in nucleosome-free regions (NFRs) flanked by the positioned nucleosomes (Kumar and Bhargava 2013). Our earlier studies had revealed a role for the terminator in regulation of chromatin transcription by the dynamics of the downstream nucleosome on pol III-transcribed tRNA genes. While Pol III is lost from the genes under repression in a gene-specific manner (Mahapatra et al. 2011; Kumar and Bhargava 2013; Shukla and Bhargava 2018) the continued TFIIIC presence on the genes (Roberts et al. 2003) protects the gene body encroachment by a nucleosome under repression. This anti-repression by TFIIIC allows chromatin modifiers to act on the locus upon activation to keep the genes nucleosome-depleted and conducive for high transcription rate. In the third step, the chromatin modifying activities like RSC and FACT regulate the terminator accessibility and the associated pol III recycling by controlling the downstream nucleosome dynamics (Mahapatra et al. 2011; Shukla et al. 2021). Apart from *SUP4*, 52 more tRNA genes in yeast show similar nucleosome dynamics near their terminator region in repressed state (Kumar and Bhargava 2013). Moreover, Pol III is found also in the region downstream of the terminator on most of the genes (Turowski et al. 2016). Although downstream termination on another terminator-like sequence is possible, most probably the downstream nucleosome on the tRNA genes (Mahapatra et al. 2011; Kumar and Bhargava 2013), blocks further pol III advancement past the terminator. Our analysis of this data showing non uniform pol III loss under starvation form the tRNA genes (Turowski et al. 2016) found that majority of the genes showing the downstream nucleosome mobility belong to the highly transcribed genes group (Shukla and Bhargava 2018).

### Terminator-dependent Pol III recycling on chromatin: A novel mechanism of Transcriptional activation

Compared to pol II, pol III is known to show much higher efficiency of transcription in vitro. On pol III-transcribed genes, the basal transcription factors TFIIIC, TFIIIB (and TFIIIA) bring promoter and terminator in close proximity probably due to the bending of DNA in the pre-initiation complex (Leveillard et al. 1991; Ferrari et al. 2004). This possibly allows direct transfer of pol III back to the TSS from the terminator, for quick reinitiation. After the first successful round, pol III can recycle 5 to 10 times faster on the same template in terminator-dependent manner (Dieci and Sentenac 1996) probably because the highly stable TFIIIB-DNA complex (Kassavetis et al. 1990) can direct multiple initiation events without letting pol III fall off the template. The polymerase recapture mechanism involving promoter bound-TFIIIB and the protein-protein interactions between PoI III and pre-initiation complex may be responsible for facilitated re-initiation on short pol III genes, while for longer templates, like *SCR1*, high re-initiation rate requires even TFIIIC (Ferrari et al. 2004). Similar to TFIIIC activity on *SNR6*, binding of transcription termination factor of RNA polymerase I (TTF1) to the upstream, promoter-proximal terminator was shown to induce remodeling and activate chromatin on the rDNA (Langst et al. 1997). No such termination factor is reported for pol III of *S. cerevisiae*. In this context, our in vitro pol III recycling experiments were performed with pure transcription components without adding any additional protein, which could affect our measurements.

The defect in terminator recognition by pol IIIΔ leads to termination in the second (T2) instead of the first (T1) terminator of *SUP4* in vitro (Landrieux et al. 2006). Since the loss of three subunits in pol IIIΔ abrogates proper termination (Landrieux et al 2006; Arimbasseri and Maraia 2013), the subsequent terminator dependent pol III recycling and reinitiation are also abolished. As discussed earlier (Shukla and Bhargava 2018), the single nucleosome dynamics in the region downstream of the terminator could have a regulatory influence on the pol III recycling and transcriptional output from the tRNA genes. Thus, terminator dependent facilitated pol III recycling turns out to be a general mechanism of high transcription activity on the pol III-transcribed chromatin in vivo; for which classical activators or enhancer-based mechanisms, prevalent for pol II-transcribed genes, are not documented. Regulation of pol II-transcribed genes generally operates from the gene upstream region on the initiation level. On the pol III-transcribed genes, the regulation comes from the gene downstream region via the transcription termination mechanism.

### Transcription regulation by the terminator-dependent cross-talk between the initiation and termination gene regions

Defective termination may generate unwanted, non-coding RNA, which may not be processed to generate the desired transcript (Orioli et al. 2011). The required accuracy and efficiency of the transcription initiation, elongation, termination and recycling phases for producing enough pol III transcripts to cater to high cellular needs; are maintained by co-operation between various factors that comprise the pol III transcription machinery, and their cognate binding sites in the DNA.

Recent advances have revealed a network of regulatory cross-talks between the three RNA polymerases (Bhargava 2021). With focus mostly on Pol I and Pol II, initiation and elongation of transcription in chromatin milieu has been extensively studied, but termination has received comparatively less attention. Terminator dependent facilitated recycling of pol III in vitro (Dieci and Sentenac 1996; Dieci et al. 2002; Cabart et al. 2008) has been documented many years back. Similarly, looping of the intervening DNA was proposed to facilitate the interaction of two proteins bound to their distant sites in space earlier (Bhargava and Chatterji 1992). Wrapping the gene body onto a nucleosome to juxtapose the initiation and termination regions of the pol III-transcribed *SNR6* may be mechanistically similar to the gene looping on a longer range found on the pol II-transcribed genes. Similar to *SNR6*, looping of the intervening DNA, which juxtaposes initiation and termination sites on mitochondrial rDNA when the termination factor binds to both of them together, results in higher transcription (Martin et al. 2005). The pol III-transcribed *SNR6* gene is atypical in its organization and behavior. The unique organizational combination of features related to both pol II and pol III transcription (Roberts et al. 1995) make yeast *SNR6* a good model gene for studying general mechanisms of eukaryotic transcription as earlier proposed for its human counterpart (Hernandez 2001). Thus, mechanisms associated with U6 gene may be extrapolated to pol II-transcribed genes as well.

Several examples in literature suggest that the gene looping mechanism of pol I and pol II transcriptional activation may be similar to the facilitated recycling of pol III on much shorter genes. These studies demonstrate that looping of DNA between termination and initiation sites is one of the ways to enhance the transcription of much longer genes (O’Sullivan et al. 2004; Ansari and Hampsey 2005; Tan-Wong et al., the RNA 3’-end cleavage and processing (West et al. 2008; West and Proudfoot 2009; Nagaike et al. 2011), transcription memory in vivo (Laine et al. and even directionality of the promoters on the bidirectional pol II promoters (Grzechnik et al. 2014). Similarly, transcription activator is proposed to promote the interaction of the initiation and termination factors on pol II-transcribed genes (El Kaderi et al. 2009), probably for facilitating the pol II recycling. On the pol III-transcribed genes, facilitated recycling was found to make the human pol III transcription refractory to repression by Maf1 in vitro (Cabart et al. 2008).

This study gives evidence in favor of a functional terminator-dependent polymerase recycling being a novel mode of transcriptional activation from chromatin templates in vivo. This facilitated recycling and consequent high rate of transcription may be a general transcription activation mechanism, which allows pol III transcription machinery to match with the high level requirement of the transcripts generated by pol III in vivo. With a wide range of processes getting influenced by the termination associated recycling of the RNA pols, it may serve as a tool to correct the defective transcription in vivo. In this context, it may be important to take note of specially pol III transcription, which had been known since long to show high levels in the cancerous cells (White 2004). A very recent review has documented several human diseases, which are manifested due to abnormal levels or functions of different tRNAs (Orellana et al. 2022). With increasing understanding of the mechanistic features of regulating pol III transcription, possibilities of exploring novel tools for better handling the pol III transcription/transcript-associated disorders improve.

## Materials and Methods

### Plasmid DNAs, yeast strains and primers

List of the plasmids, yeast strains and primers used in this study may be found in the Supplemental Tables S1,2,3. Plasmids carrying the wild type (pCS6; Brow and Guthrie 1990) and modified *SNR6* gene (pU6LNS; Pazin et al. 1997) were previously described (Shivaswamy and Bhargava 2006). Terminator was inserted in pU6LNS (Supplemental Table S1, Fig. S1A) to give either pU6LR (termination at +163, downstream) or pU6LNST (termination at +116). Terminator of the gene in pCS6 and pU6Maxi (pB6M; Burnol et al. 1993b) was disrupted to give the plasmids pU6dT and pU6dTMaxi respectively. *SNR6* gene with disrupted terminator was cloned in the plasmid pRS306 (Sikorski and Hieter 1989) and plasmid shuffle method was used to create the strain YPB110 with terminator disruption at the gene locus. Strains YPB17 (used as control strain) and YPB110 (both in W3031a background) carry wild type U6 gene in pRS413. The largest pol III subunit C160 (in W3031a cells) and the second largest subunit of pol III, C128 in the wild type strain YPH500 and the strain C37HAΔCt (Landrieux et al. 2006) were 3X-myc tagged using PCR Tool Box (Euroscarf) according to Janke et al. (2004).

### Chromatin assembly and transcription in vitro

Chromatin was assembled on plasmid DNAs using S-190 extract of Drosophila embryos (Pazin et al. 1997), which does not have any activity that supports the naked DNA transcription by pol III in vitro (Shivaswamy et al. 2004). Templates with a ladder of at least 6 discernable nucleosomes were used for all the experiments. In vitro transcription on naked DNA and chromatin templates using pure yeast TFIIIC, recombinant TFIIIB and pure yeast pol III and quantifications were as described previously (Shivaswamy et al. 2004). The mutant form of pol III lacking subunits C11, C37 and C53; Pol IIIΔ was purified from the C37HAΔCt cells according to Landrieux et al. (2006). The transcript was visualized by primer extension method using AMV reverse transcriptase. Transcript yield in each lane was normalized with respect to the recovery marker and fold activation of transcription in the presence or absence of TFIIIC was calculated as ratio of normalized transcript yields in each case. Average from at least three independent experiments with scatter is given in the graphs.

Details of ATP-depletion experiment were as described earlier (Shivaswamy et al. 2004; Shivaswamy and Bhargava, 2006). Replenishing includes adding back 3 mM ATP and creatine phosphate to original levels while all four NTPs were added in much lower amounts (0.5 mM) for transcription in every condition. Fold activation was calculated as ratio of transcript yield with or without TFIIIC addition as described before.

### Chromatin structure analysis

Indirect end labeling (IEL) analysis (Vinayachandran et al. 2009) was used to analyze chromatin structure at longer range and MNase footprinting was used to analyze chromatin structure at closer range. MNase digestions for IEL and footprinting analysis were carried out as described previously (Shivaswamy et al. 2004) for pU6LNS and pU6LR.

### Chromatin immunoprecipitation (ChIP) and RNA analysis in vivo

Pol III occupancy was measured at gene loci using strains having Myc-tagged C160 or C128 subunits and Anti-myc antibody (Santacruz). Chromatin was MNase digested to monosome level for ChIP samples preparation. TELVIR was used as normalizer. ChIP and Real time PCR, RNA extraction and quantification were performed as described (Arimbasseri and Bhargava 2008; Mahapatra et al. 2011), and repeated at least three times for each experiment. The ChIP data were analyzed essentially as described by Aparicio et al. (2004), using ΔΔCt method. Statistical significance of the differences was calculated by applying two-tailed Student’s t test.

### Data availability

All data supporting the findings of this study are available here and in the Supplemental Material.

## Supplemental data

The supplemental data include three Tables and five Figures.

## Acknowledgements

We thank Michel Riva for Pol IIIΔ strain and Christine Conesa for U6 maxi plasmids. Financial help from CSIR, Govt. of India is acknowledged. VV and SS were recipients of CSIR and AS was recipient of Indian Council of Medical Research (ICMR) Senior Research fellowship.

## Author Contributions

VV designed, performed most of the experiments and analysed the results, SS did the footprintings and chromatin structure analyses, AS performed the ChIP, RT-qPCRs and analysed the data. VV designed, created and NK created the termination mutations; both constructed the modified yeast strains. PB conceived the study, designed the experiments and wrote the paper.

