## Supplemental Tables for "Terminator-dependent facilitated recycling of RNA polymerase III couples transcriptional activation and chromatin remodeling in vivo"

for the manuscript

**Contents** of this File

I. Supplementary Tables S1, S2 and S3

### I Supplemental Tables

#### Table S1 List of the plasmids used in the study

| Plasmid | Reference | Comment/Destails of the plasmid |
|---|---|---|
| pCS6 | Brow and Guthrie 1990 | The plasmid contains the 432 bp DNA from the positions -120 to +312 bp of yeast genomic DNA harboring the *SNR6* gene. |
| pU6LNS | Pazin et al. 1997; Shivaswamy and Bhargava 2006 | The plasmid was derived from pCS6. The gene terminator sequence has been deleted and a 50 bp synthetic sequence was put between the box A and box B . |
| pU6LR | This study Fig. S1A | The U6 terminator sequence was inserted in pU6LNS at two positions. One downstream of B box for transcription termination at +163 bp and the other upstream of U6TATA box to terminate a leftward transcription at -65 bp position. |
| pU6LNST | This study Fig. 3A | The plasmid was made by inserting the U6 terminator sequence in pU6NLS to give transcription termination at +116 bp position. |
| pU6dT | This study Fig. 3A | The plasmid was made by disrupting the terminator stretch of ten 'T' residues in pCS6 to a sequence TTTTAAATTT, employing the site-directed mutagenesis. |
| pRS306 | Sikorski and Hieter (1989) | A gift from Trisha Davis, University of Washington. |
| pRS413 | Sikorski and Hieter (1989) | A gift from K Natarajan, Jawahar Lal Nehru University, New Delhi. |
| pU6M | pB6M, Burnol et al. (1993b) | The plasmid carries the maxi version of the wild type SNR6 gene which gives a transcript of the size 232 nt. |
| pU6MdT | This study | Same as pU6M but with the disrupted terminator sequence TTTTAAATTT. |

#### Table S2 List of the yeast strains used in this study

| | | | |
|---|---|---|---|
| 1 | W3031a | MATa, ade2-1, his3-11, -15, leu2-3, -112, trp1-1, ura3-1, can1-100 | Lab stock |
| 2 | C37HAΔCt (Pol IIIΔ) | MATα ura3-52 lys2-801_amber ade2-101_ochre trp1-Δ63 his3-Δ200 leu2-Δ1 rpc37Δ18-3HA::kanMX6 *RPC160: HIS3* | Michel Riva |
| 3 | YPH500 | MATα ura3-52 lys2-801_amber ade2-101_ochre trp1-Δ63 his3-Δ200 leu2-Δ1 | Magdalena Boguta |
| 4 | yPB31 | MATα ura3-52 lys2-801_amber ade2-101_ochre trp1-Δ63 his3-Δ200 leu2-Δ1 RPC128-3xmyc::TRP1 | This study |
| 5 | yPB32 | MATα ura3-52 lys2-801_amber ade2-101_ochre trp1-Δ63 his3-Δ200 leu2-Δ1 rpc37Δ18-3HA::kanMX6,RPC128-3xmyc::TRP1 | This study |
| 6 | yPB17 | MATa, ade2-1, -15, leu2-3, -112, trp1-1, ura3-1, can1-100, his3-11, *pRS413(HIS3,* CEN6, *ARSH4, SNR6)* | This study |
| 7 | yPB110 | MATa, ade2-1, -15, leu2-3, -112, trp1-1, ura3-1, can1-100, his3-11, *SNR6-dT::URA3 pRS413(HIS3,* CEN6, *ARSH4, SNR6)* | This study |

**Table S3 Primers used in the study**

| Primer Name | Sequence 5'--------3' | Used for |
|---|---|---|
| RT Both (P2/ Gene) | TCTCTTTGTAAAACGGTTCATCCT | PE; Fig. 1A, 3B, 4A, 4D, Fig. S3, S4B,C |
| U6maxiRT (P1) | GGCTGCAGGAATTCGATATCAAGC | PE; Fig. 3D; Fig. S3A,B |
| RT dwn (DS) | GTACGCTGGGTATCTTCAAATGTAC | PE; Fig. S3 C,D; S4C |
| SNR14 (U4) | GCGAACACCGAATTGACCATG | PE; Fig. 4D, Fig. S4C |
| U4298 F | CACGGGAAATACGCATATCAGTGA | RTPCR, RNA; Fig. S4C |
| U4299 R | CCGAATTGACCATGAGGAGACG | RTPCR, RNA; Fig. S4C |
| Upstream F | GTCATCTTCCTGGACCTCATG | ChIP-RTPCR, Fig. 3H, Fig. 4F |
| Upstream R | GCAATGAAACTCTAAAGTATCATCGAT TCAG | ChIP-RTPCR, Fig. 3H, Fig. 4F |
| TATA Box F | CGATGATACTTTAGAGTTTCATTGC | ChIP-RTPCR, Fig. 3H, Fig. 4F |
| TATA box Rev | CTTCGCGAACACATAGTTGC | ChIP-RTPCR, Fig. 3H, Fig. 4F |
| A-B Box F | GTTCCCCTGCATAAGGATGAACCG | ChIP-RTPCR, Fig. 3H, Fig. 4F |
| A-B Box R | GGAAGATAAAGATACACTGCTG | ChIP-RTPCR, Fig. 3H, Fig. 4F |
| U6down for | GTACTTATGTGCTTTATGAATGTG | ChIP-RTPCR, Fig. 3H, Fig. 4F |
| U6down rev | CCTTTCTCTTCTGTTTGACA | ChIP-RTPCR, Fig. 3H, Fig. 4F |
| Tel VIR F | GCGTAACAAAGCCATAATGCCTCC | ChIP-RTPCR, Fig. 3H, Fig. 4F |
| Tel VIR R | CTCGTTAGGATCACGTTCGAATCC | ChIP-RTPCR, Fig. 3H, Fig. 4F |
| SUP4 Up F | TGGTCATGATGTCGCTATTTCT | ChIP-RTPCR, Fig. 4E |
| SUP4 Up R | CTAAAGCTAAGGGATGGAAGAGA | ChIP-RTPCR, Fig. 4E |
| SUP4 Gene F | CTCTCGGTAGCCAAGTTGGTTTAAGGC | ChIP-RTPCR, Fig. 4E |
| SUP4Gene R | CCCGGGGGCGAGTCGAACGCCCGA | ChIP-RTPCR, Fig. 4E |
| SUP4 Down F | CCCGGGAGATTTTTTTGTTT | ChIP-RTPCR, Fig. 4E |
| SUP4 Down R | AAAAGAGGCTACAAGAGTTCGTTAAT | ChIP-RTPCR, Fig. 4E |
| SUP4 IN | ATCTCAAGATTTCGTAGTGA | PE intron-specific; Fig. 4D |

PE stands for primer extension, RT for Real Time.
