## Supplemental Figures for "Terminator-dependent facilitated recycling of RNA polymerase III couples transcriptional activation and chromatin remodeling in vivo"

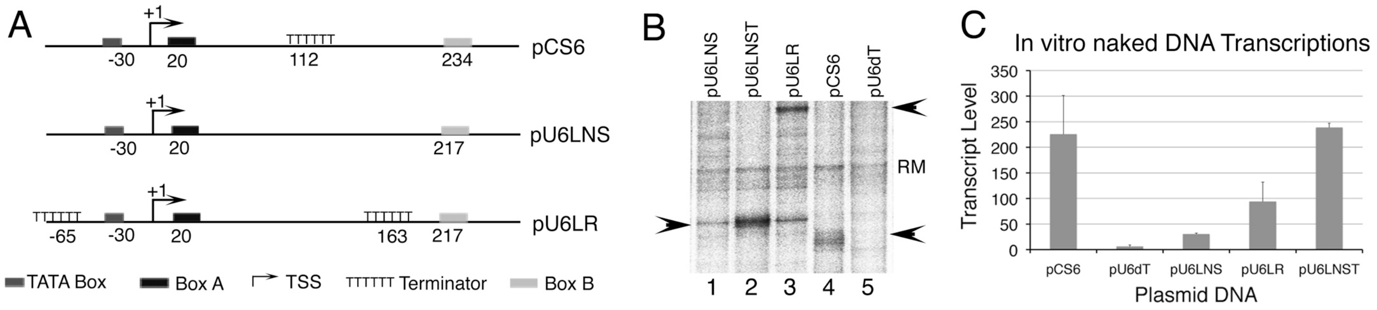


**Figure S1. Requirement of terminator for transcription of the yeast U6 snRNA gene.** (A) Schematic representation of the wild type *SNR6* gene in pCS6 and its modified forms in pU6LNS and pU6LR (Supplemental Table S1). Key to all symbols used is given in the bottom of the panel. Scheme for pU6dT and pU6LNST may be found in the Figure 3A. Transcription was followed on naked DNA templates by UTP incorporation method. (B) In vitro transcription using pure pol III transcription machinery. Transcription on naked plasmid DNA templates using pure yeast TFIIIC, recombinant TFIIIB subunits TBP, Brf1 and Bdp1 and pure yeast pol III was carried out using $\alpha$-[$^{32}$P]-UTP to visualize the transcript. Arrows mark the right size transcripts from pU6LNST, pU6LR and pCS6 in the lanes 2, 3 and 4 respectively. In the absence of terminator in pU6dT and pU6LNS, no distinct transcript of expected size is seen. In the lanes 1-3, some background bands from the three LNS plasmids are seen overlapping the position in gel for the right size transcript from pU6LNST (lane 2, band marked with arrow). These are possibly because of the A-T rich synthetic sequence inserted in the plasmid while deleting the U6 terminator from pCS6 for construction of the plasmid pU6LNS.
(C) Yield of transcript in each case was normalized against recovery marker. Average from three independent experiments with scatter is shown.

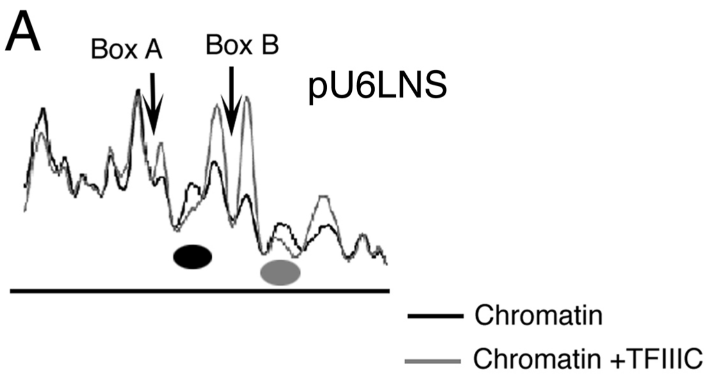


**Figure S2. Binding of TFIIIC leads to nucleosome positioning on the body of the modified U6 genes.**

Indirect End-labeling (IEL) analysis of chromatin structure showing profile matchings of lanes from the gel in the Figure 2A on the modified U6 genes (A) without (pU6LNS, upper panel) and (B) misplaced (pU6LR, lower panel) terminator. Vertical arrows mark the positions of the boxes A and B while ellipses mark the positioned nucleosomes. Comparison of the aligned profiles of similarly digested chromatin shows that similar to the wild type gene, a chromatin remodeling in the presence of TFIIIC results in positioning of a nucleosome (a clear protected region) between the boxes A and B of both the modified genes. A protection downstream of box B (black ellipse) is also seen.

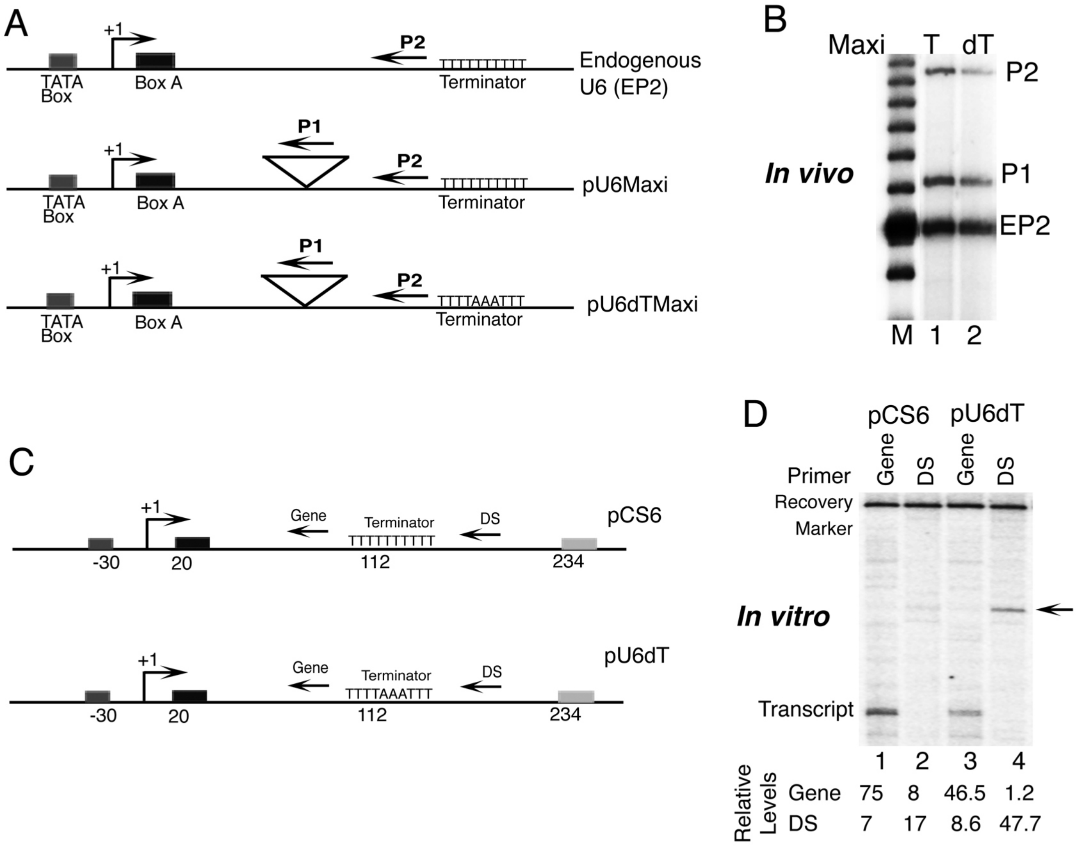


**Figure S3. Transcription from the *SNR6* with disrupted terminator in vivo and in vitro**
(A) Cartoons show the scheme of the U6 maxi genes carrying normal (pU6Maxi) or disrupted (pU6dTMaxi) terminator as compared to the gene at the endogenous *SNR6* locus. Triangle shows the insert in the Maxi gene. Positions of the primer extension probes P1 and P2 complementary to top strand are shown; P2 probes RNA from endogenous locus also (EP2).
(B) A gel showing primer extension products with P1 and P2 from total RNA isolated from cells carrying pU6Maxi (T) or pU6dTMaxi (dT). EP2 shows endogenous U6 product with P2. M shows a 10 bp DNA ladder used as molecular size marker.
(C) Position of the reverse transcription primers for visualizing the transcripts from the *SNR6* gene. Cartoon shows the positions of the primers with respect to the terminator, used for primer extension in panel D. Gene and DS (downstream) are the primers hybridizing to coding region and its downstream respectively.
(D) Primer extension products from the RNA synthesized by wild type Pol III in the presence (pCS6) or absence (pU6dT) of the terminator in vitro. Arrow on the right hand side of the gel points to the primer extension product (168 nt) using DS primer from naked DNA transcription of pU6dT. Relative RNA levels of each type of transcript after normalization with recovery marker are given in the bottom of the gel. With background level of 7-8, negligible transcript from downstream of pCS6 can be seen while a lower level transcript from the gene region, and ~ 3 fold more transcription from downstream region of pU6dT is seen.

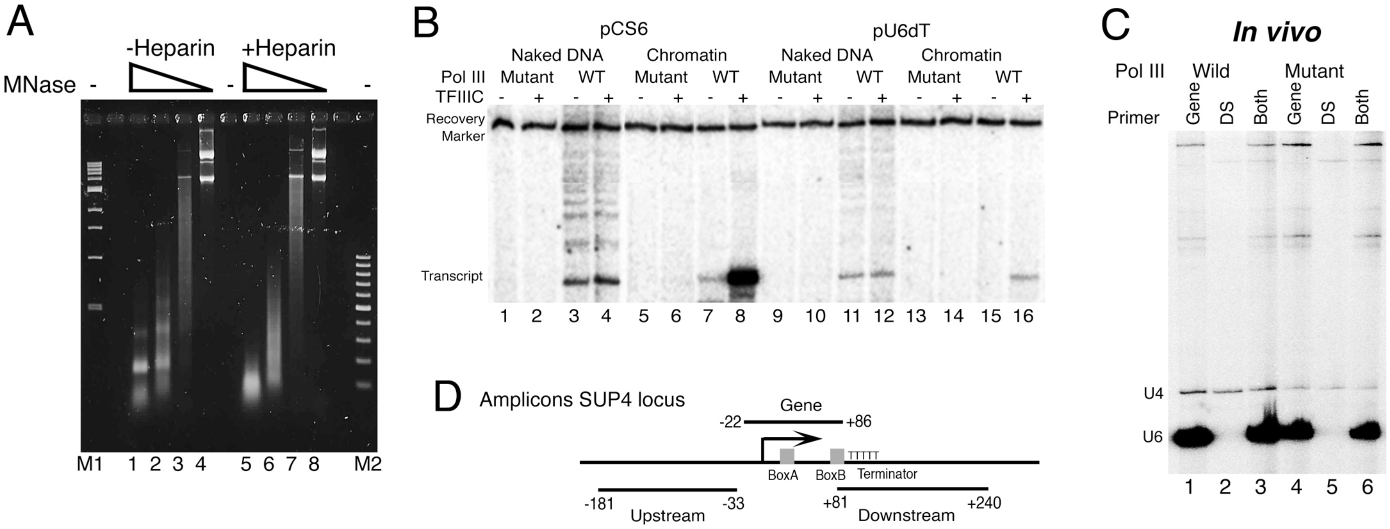


**Figure S4.** Mutant pol III transcribes poorly in vitro and in vivo
(A) Effect of heparin on chromatin structure. Chromatin assembled on the plasmid using S-190 extract. After 4.5 hr of assembly, 300 μg/ml heparin, same as the level required in single round transcription experiments, was added and incubated for further 15 mins., before subjecting to MNase digestion for ladder assay. Disruption of ladder and presence of mononucleosomal size product as major species shows heparin results in loss of proper chromatin structure.
(B) Mutant pol III transcribes poorly in vitro and in vivo.
Comparison of transcription by the wild type (WT) and mutant pol III. Representative gel showing transcription from chromatin assembled on the plasmids pU6dT and pCS6 in vitro, with or without TFIIIC addition. Positions of the recovery marker and the transcript are marked.
(C) No downstream transcription by mutant pol III is seen in vivo.
Primer extension products from RNA isolated from the cells carrying wild type (YPH500) or mutant (C37HAΔCt) Pol III. U4 was used as normalizer in each lane. Gene and DS (downstream) are the primers hybridizing to coding region and downstream of the gene respectively. Gene probes for U6 in lanes 1, 3, 4 and 6 while no right size extension product with DS primer is seen in the lanes 2, 3, 5 and 6.
(D) Cartoon showing positions of the amplicons for measuring Pol III enrichment by ChIP-Real Time PCR method as described under methods on the *SUP4* gene locus. Downstream amplicon has an overlap in terminator region with the Gene amplicon.

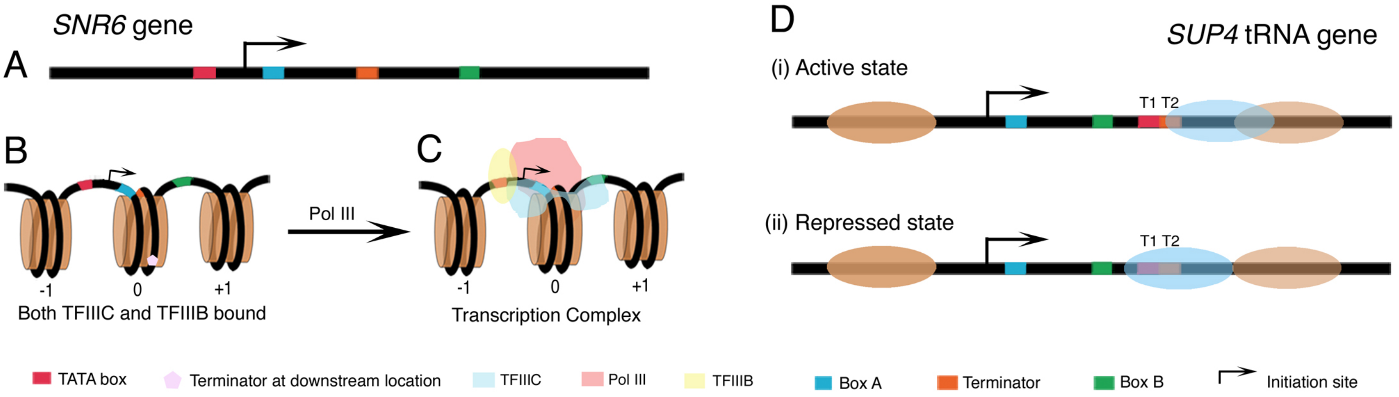


**Figure S5. Two representative ways of gene regulation by terminator-dependent facilitated recycling of pol III on the chromatin.**

Steric layout of pol III transcription machinery over the *SNR6* (A-C) and *SUP4* tRNA (D) genes in nucleosomal context. Cartoons are color coded. Black bar represents DNA.

(A) Linear representation of the *SNR6* gene. Terminator of *SNR6* is located between the two promoter elements in linear arrangement. A positioned nucleosome on the gene region was proposed to facilitate TFIIIC binding by bringing its distant binding sites, A and B boxes close in space and enhance transcription from chromatin (Shivaswamy et al. 2004).

(B) Arrangement of positioned nucleosomes (brown cylinders) on the gene locus once TFIIIC and TFIIIB (not shown for the sake of clarity) are bound. The -1 nucleosome is found upstream of -70 bp position, while positioning of a nucleosome on the gene body brings the Boxes A, B and terminator close in space such that terminator is oriented away from the boxes but close to the transcription initiation region. Normal terminator is close to the dyad axis while misplaced terminator is found on the opposite side of the dyad axis.

(C) Steric layout of pol III transcription machinery over the *SNR6* gene, with respect to the nucleosome between the boxes A and B. The initiation complex is assembled in the NFR region, whereas the presence of nucleosome on the gene body helps keep the whole assembly very compact. Halfway, near the dyad axis, terminator allows pol III to disengage from the template and move directly to initiation site again, making facilitated recycling possible in the presence of bound TFIIIC.

(D) Terminator-directed facilitated recycling of pol III on tRNA genes is controlled by the downstream nucleosome dynamics. Representative example of *SUP4* gene has two transcription terminators T1 and T2 as shown. Blue oval represents the downstream dynamic nucleosome. (i) In active state, pol III is directly transferred from terminator to initiation region for recycling (Dieci and Sentenac 2003). A fuzzy downstream nucleosome supports the pol III recycling from T1 by keeping the terminator accessible, which allows high transcription rate. (ii) In repressed state, a positioned nucleosome blocks both T1 and T2, bringing transcription to a halt, though the gene may be still occupied by its transcription factors.
